# Morphine self-administration induces region-specific brain volume changes and microglial phenotypic alterations without affecting neuronal density in male Wistar rats

**DOI:** 10.1101/2025.02.20.639378

**Authors:** Ana Débora Elizarrarás-Herrera, David Medina-Sánchez, Mariana Stefania Serrano-Ramírez, Diego Angeles-Valdez, Luis A. Trujillo-Villarreal, María Antonieta Carbajo-Mata, César J. Carranza-Aguilar, Eduardo A. Garza-Villarreal

## Abstract

Addiction to opioids, including morphine, is a major public health crisis in the U.S. It has been associated with brain volume changes in reward-related regions, neuronal death, and neuroinflammation. However, the link between structural changes and neuroinflammation is not well understood. In this study, we used operant conditioning to induce morphine self-administration in rats and examined brain volume and cellular changes, focusing on microglial phenotypes. Male Wistar rats were conditioned to morphine self-administration (0.01 mg/kg) for 20 days under a fixed-ratio 1 schedule. In vivo structural Magnetic Resonance Imaging (MRI) scans were conducted at the beginning and end of self-administration. Brains were stained for Iba1 and NeuN proteins, and confocal images were analyzed for cell counts and microglial morphology. We used Deformation-Based Morphometry for MRI volume analysis and Principal Component Analysis with K-means clustering for microglial phenotyping. Our results showed that morphine self-administration led to volume changes in addiction-related brain regions, including increased globus pallidus and decreased insular cortex volume. Additionally, morphine caused widespread neuroinflammation, evidenced by elevated microglial density in the caudate-putamen, dentate gyrus, globus pallidus, and insular cortex, without affecting neuron counts. Finally, we observed region-specific variations in microglial phenotypes, suggesting region-specific neuroinflammatory roles. In conclusion, our study shows that morphine self-administration induces structural and microglial changes in addiction-related brain regions without neuronal loss, highlighting the role of neuroinflammation in opioid-induced adaptations. The variability in microglial phenotypes underscores their complexity, emphasizing the need to study their progression in addiction and their potential as therapeutic targets.

## 1. Introduction

Morphine is an opioid widely used in the treatment of acute and chronic pain (Murphy, Bechmann, and Barrett 2024). However, it is also known for its ability to induce brain changes that result in physical dependence and addiction, even at low doses and after short-term use (Strang et al. 2020; Lin et al. 2016). Non-desired effects of morphine are related with several neuroadaptations at the central nervous system (SNC) level, including macrostructural alterations (Upadhyay et al. 2010) and neuroinflammation (Carranza-Aguilar, Rivera-García, and Cruz 2022; Cahill and Taylor 2017).

Brain volumetric changes have been reported as a result of prescription opioid treatments and in opioid abusers (Younger et al. 2011; Borne et al. 2005). In humans, chronic opioid use leads to a reduction in the volume of brain regions involved in decision-making, such as the prefrontal cortex (Lyoo et al. 2006), as well as regions responsible for emotional regulation, like the amygdala (Lin et al. 2016; Upadhyay et al. 2010), and memory, including the hippocampus (Younger et al. 2011). Similar reductions and some increases have been observed in the hippocampus and nucleus accumbens in animal models of opioid self-administration (Taylor et al. 2023; Cannella et al. 2024). These effects potentially modify specific functions such as rewarding and memory, contributing to the addictive potential of morphine. Based on particular structural alterations, opioid-induced volumetric changes could be related to different mechanisms such as cell death, proliferation, recruitment of immune cells, neurogenesis or neuroimmune responses (Datta et al. 2017; Cannella et al. 2024).

Neuroinflammation is an immune response in the central nervous system (CNS) primarily mediated by microglia, specialized cells that contribute to maintaining homeostasis (Schetters et al. 2017; Sierra, Paolicelli, and Kettenmann 2019). Depending on the nature of the disturbance, extent of damage, receptors involved, types of cells affected, and duration of stimuli, microglia can exhibit diverse responses and morphologies (Woodburn, Bollinger, and Wohleb 2021). Generally, microglia display various phenotypes that reflect their functional roles in the brain: ramified microglia have small cell bodies with numerous long processes crucial for environmental surveillance; hyper-ramified microglia show extensive branching associated with pro-inflammatory activity, potentially modulating neuroinflammatory responses; hypertrophic (bushy) microglia possess enlarged bodies with shorter, stubby branches, indicating phagocytic activity for tissue clearance; ameboid microglia feature rounded cell bodies and retracted branches, primarily functioning in increased phagocytosis of extracellular proteins and dying cells; and rod-like microglia have elongated bodies and fewer branches, detecting axons and neurons to provide neuroprotection (T. R. F. Green and Rowe 2024).

The diversity of microglial responses to morphine may be influenced by several factors, including the receptors involved, such as opioid receptors or toll-like receptors, the type of neuroinflammatory response, and the specific brain region affected (King’uyu et al. 2023; Y. L. Chen, Law, and Loh 2006). It has been reported that microglial activation in the hippocampus produces pro-inflammatory cytokine release, resulting in altered synaptic plasticity and cognitive dysfunction (Cornell et al. 2022). In the prefrontal cortex, it is linked to changes in mood and behavior (Guo et al. 2023), whereas in the spinal cord, microglia contribute to neuropathic pain by releasing neurotrophic factors (G. Chen et al. 2018). Additionally, in regions critical for addiction, morphine-induced microglial activation can enhance reward and addictive behaviors, highlighting the region-specific nature of microglial activity (Lacagnina, Rivera, and Bilbo 2017). The role of microglia in brain regions involved in the intoxicating effects of morphine, particularly those with high opioid receptor expression, such as the caudate-putamen (CPu), globus pallidus (GP), insular cortex (InsCx), and specific regions of the hippocampus, has not been completely elucidated (J. M. Green, Sundman, and Chou 2022; Carranza-Aguilar et al. 2022; Blackwood and Cadet 2021). In addition, while there are some insights into the structural consequences of morphine in the brain, the specific mechanism involved and the reason why certain regions are more susceptible to changes than others remain unclear.

Understanding the neurobiological mechanisms underlying morphine addiction is crucial for the development of effective treatment strategies. Moreover, the characterization of different phenotypes of microglia could serve as a biological marker for the mechanisms that morphine and other substances can produce. Here, we propose that morphine self-administration may induce alterations in the volume of brain regions associated with addiction, and suggest that these changes could be linked to diverse regional susceptibility of the brain to change microglial state. Therefore, the aims of this study were to: a) investigate the macrostructural brain changes induced by morphine self-administration using magnetic resonance imaging, b) examine whether morphine self-administration alters neurons and microglia counts in brain regions associated with the development of addiction, and if these are related to brain volume changes, and c) determine whether different reactive microglia phenotypes are expressed in brain regions involved in morphine self-administration behavior.

## 2. Materials and methods

### 2.1. Animals

Male Wistar rats were obtained from the Institute of Neurobiology’s vivarium and individually housed in polypropylene cages, under inverted light/dark cycle (12/12 h; lights on at 08:00 p.m.), with controlled temperature (22 – 24 °C) and access to food and water *ad libitum*. All methods were approved by the Bioethics Committee of the Institute of Neurobiology (protocol 113.A). They were carried out in accordance with the regulations established by the Mexican Official Standard for the care and handling of animals (1999), complied with the National Institutes of Health guidelines (NIH 2011) and followed the ARRIVE guidelines for animal research (Kilkenny et al. 2010).

### 2.2. Drugs

Morphine sulfate was administered intravenously at 0.01 mg/kg/infusion. Isoflurane (Sofloran) was our anesthetic of choice both for surgical procedures as well as the magnetic resonance imaging procedures (5% for induction and 1.5% for maintenance). For postoperative care of the animals, we used the non-steroidal anti-inflammatory Meloxicam (1 mg/kg), and Enrofloxacin (10 mg/kg) for prophylactic antibiotic coverage. In order to maintain our catheter permeability and ensure long-term patency, we opted for a flushing solution consisting of the antibiotic Ticarcillin (10 U/ml) combined with the anticoagulant Heparin (66.67 mg/ml). An anesthetic cocktail of Ketamine (15 mg/ml) and Midazolam (0.75 mg/ml) was used for mild sedation to test the patency of the catheter. All drugs were from Sigma-Aldrich (Toluca, Mexico).

### 2.3. Experimental Design

A total of 12 male Wistar rats were acquired on postnatal day (P) 37 and habituated to handling (5 min/day) for 5 days to reduce stress from manipulation. Surgeries for intrajugular cannulation with button ports were performed on P42, followed by a 2-week recovery period. On P57, structural magnetic resonance imaging (MRI; T1) was performed. Over the next three days, all rats were habituated to operant conditioning chambers, and on the fourth day, a successive lever approximation protocol (shaping) was performed. Next, animals were randomly assigned to two groups: five rats were exposed to self-administration of 0.01 mg/kg/100µl morphine (Mor group), and six were exposed to vehicle (0.9% sodium chloride) infusion (Ctrl group). A rat in the Mor group was discarded due to the development of infection after surgery (see Surgery section). On day P62, we initiated a fixed ratio 1 (FR1) self-administration schedule, where each correct response was rewarded, and continued for 20 sessions. Four hours after the last session (P81), an additional structural MRI (T2) was performed. Finally, rats were perfused, and their brains were extracted and cryoprotected until immunofluorescence processing (see Figure 1 for details).

**Figure 1.**
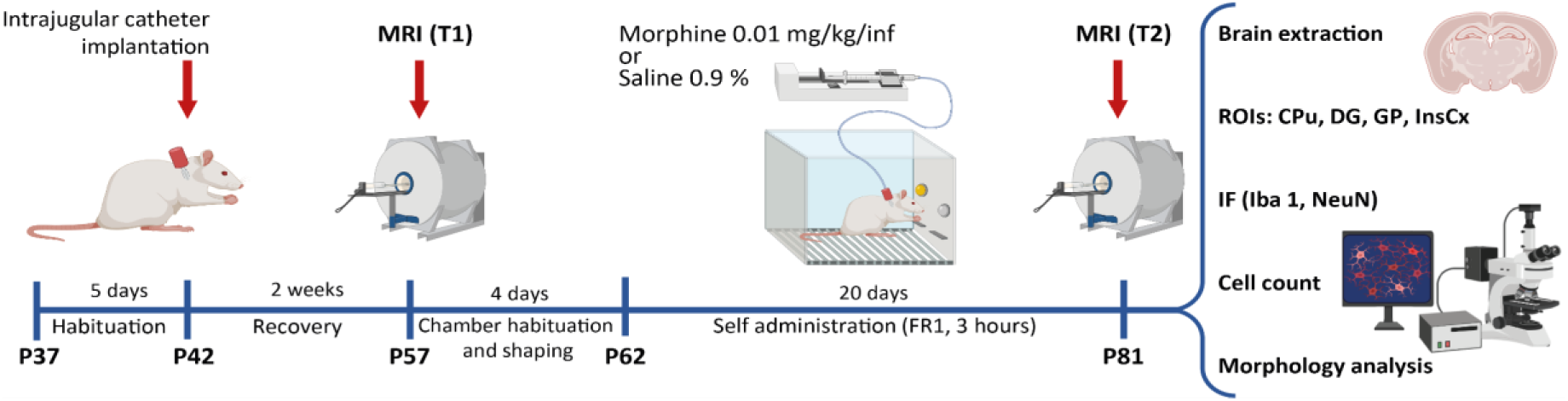
Experimental design. Rats were acquired on postnatal day (P) 37 and received habituation to handling until P42. A catheter was surgically implanted in the jugular vein, followed by a recovery period that lasted until P57. In order to produce familiarization to the operant conditioning chambers, rats were introduced to chambers (1h/day) for three days and, on the fourth day, underwent a successive lever approximations protocol (shaping). On P62, both the morphine (Mor) as the saline (Ctrl) group were introduced for three hours to the operant chambers to produce self-administration behavior under a FR1 schedule. Structural magnetic resonance imaging (MRI) scans were obtained on P57 and P81. At the end of the protocol, brains were extracted and regions of interest (ROIs) were located (CPu: Caudate Putamen, DG: Dentate Gyrus, GP: Globus Pallidus, InsCx: Insular Cortex) to stain and analyze the number of neuronal (NeuN) and microglial (Iba1) cells and morphology of microglia.

#### Surgery and post-operative care

Rats weighing approximately 200 gr were anesthetized with a 5% isoflurane for induction and 1.5% for maintenance in oxygen-enriched air. Anesthesia was confirmed by the lack of a pedal reflex response to pressure on the metatarsals of the rat. The surgical site was clipped and aseptically prepped with Chlorhexidine scrub and 70% alcohol, and the right jugular vein was exposed. A 22-gauge 3Fr catheter, made of polyurethane (C30PU-RJV1301), was inserted in the vein, secured with sutures, and tunneled under the skin to the back of the rat using a trocar ending at a small superficial incision located at the level of the shoulder blades. Once the end of the catheter was exposed, it was connected to a non-magnetic vascular access button (VAB95BS) and incisions were closed with nylon sutures. To protect the access port, the vascular access button was covered with a removable aluminum cap (catheter and accessories were purchased from Instech Laboratories, Inc.). The first five days following surgery, enrofloxacin (10 mg/kg) diluted 1:1 with saline and meloxicam (0.2 mg/kg for the loading dose, followed by 0.1 mg/kg), were administered subcutaneously for the first five days post-surgery. Additionally, the catheter was flushed daily with ticarcillin antibiotic (66.67 mg/ml) and 100 µl heparinized saline (20 IU/ml) until full recovery. Rats completely recovered within 10 to 12 days after surgery. Thereafter, catheters were flushed daily with heparinized saline until the end of the experiment. We made certain to utilize positive pressure techniques in order to ensure extended patency of the catheters. Catheter permeability was tested by flushing it with 100 µl of a solution containing 15 mg/ml ketamine and 0.75 mg/ml midazolam which produces the loss of righting reflex within 3 seconds of infusion. Rats that did not exhibit this response, or that experienced catheter occlusion, infection, or pain (n = 1) were euthanized and excluded from the analysis.

### 2.4. Self-administration protocol

#### Apparatus

The operant conditioning self-administration system was (Med Associates Inc.) comprised of twelve extra tall chambers (MED-007-CT-D1), each equipped with two retractable levers positioned 5 cm above the grid floor, and yellow cue lights located 5 cm above the left and right levers. Chambers were placed in sound-attenuating boxes (ENV-018V) to reduce external noise and were connected to infusion pumps (PHM-210) for substance administration through a 25-gauge drug delivery line.

#### Habituation and shaping

To familiarize the rats with the chambers, they were introduced for 1 hour each day over three days. On the fourth day, rats were randomly assigned to the morphine group (n = 5) or the saline group (n = 6) and underwent a successive lever approximation protocol to help them realize that pressing the lever had a consequence. To do this, rats were individually placed in the chamber with the catheter connected to the vascular access port. The right retractable lever was exposed and the rat was allowed 5 minutes of free exploration. Next, morphine (0.01 mg/ml/20µl) or saline was delivered each time the animal met one of the following criteria: 1) approaching the lever, 2) sniffing it, 3) placing a paw on it, and 4) pressing it. The substance was released 5 times for each criterion, in the order presented. The shaping session concluded after one hour or upon meeting all criteria, after which the rats were returned to their cages.

#### Self-administration

One day after the shaping protocol, both morphine and control rats were individually placed in the chambers and connected to infusion pumps. Each session lasted 3 hours and was conducted for a total of 20 sessions, starting at 10:00 AM during the dark phase of the cycle. A fixed-ratio 1 (FR1) schedule was used for substance delivery, where pressing the left lever activated the pump to infuse either saline or morphine (0.01 mg/kg/100 µl) for 5 seconds at a rate of 20 µl/sec; pressing the right lever had no effect. An illuminated cue light above the left lever indicated the availability of the substance. After lever pressing, the light was turned off for 30 seconds, during which lever presses did not result in further infusion. Infusion counts and the number of active (left) and inactive (right) lever presses were recorded each time the levers were pressed, during the 5-second infusion period, and during the 30-second timeout.

#### Motivation and perseverance measurement

Given that compulsive seeking of drugs is an indicator of addiction-like behavior, data from the last five sessions were used to obtain motivation and perseverance scores (Le et al. 2017; Bock et al. 2013). Motivation was measured by counting the total number of correct lever presses until the last infusion, indicating the effort to obtain the maximum amount of morphine per session. The perseverance value was calculated as the mean correct lever presses during the timeout and infusion periods, during which morphine was not available, reflecting the difficulty of stopping drug use. A combined behavior score was then calculated for each rat by adding the z-scores of perseverance and motivation values. By using z-scores, which have a mean of zero and a standard deviation of one, we ensured equal contribution of each behavior and eliminated inherent differences in the variance of each measurement.

### 2.5. Magnetic Resonance Imaging acquisition

The *in vivo* MRI acquisition was conducted at the National Laboratory for Magnetic Resonance Imaging using a 7 T MRI scanner (Bruker Pharmascan 70/16 US) with a 2x2 array surface rat coil and the Paravision 6.0.1 console (Bruker, Ettlingen, Germany). All rats were anesthetized during scanning sessions using a protocol adapted from Sirmpilatze et al. (2019). A single bolus of 0.012 mg/kg of dexmedetomidine was administered subcutaneously, then Isoflurane at 5% in an oxygen/air mixture (1:1) was used for induction and maintained at ∼1%. Body temperature was maintained at 38 °C by warm water circulation. Respiration rate and blood oxygen saturation was monitored with an MRI-compatible pulse oximeter and the software PC-SAM (v8.02). Structural MRI was acquired with a T2-weighted 3D FLASH sequence with following parameters: TR = 30.76 ms, TE = 5 ms, flip angle = 10°, slice thickness = 25.6 mm, FOV = 28.2 x 19 x 25.6 mm, isometric voxel = 160 µm, with two repetitions and sagittal as primary slice direction. For image preprocessing, all images were converted to NIfTI format using the brkraw tool v0.3.7 (https://brkraw.github.io/). All MRI images were preprocessed using an in-house pipeline incorporating the MINC-toolkit and ANTs packages. The structural preprocessing included intensity normalization, image centering, denoising, and registration using LSQ6 alignment. (https://github.com/psilantrolab/Documentation/wiki/Preprocessing-Rat-Structural-in-vivo). Finally, we applied Deformation Based Morphometry (DBM) employing a Two-Level DBM approach (https://github.com/CoBrALab/optimized_antsMultivariateTemplateConstruction) with the SIGMA anatomical template v. 1.2.1 (Barrière et al. 2019) to derive relative Jacobian determinants for each subject and session as a measure of volume difference at each voxel in the image relative to the average.

### 2.6. Immunofluorescence

Twenty-four hours after the last self-administration session, rats were anesthetized with pentobarbital (65 mg/kg, i.p.) and transcardially perfused with 200 ml saline solution, followed by 200 ml of 4% paraformaldehyde (PFA) in 0.1 M phosphate-buffered saline (PBS; pH = 7.4). The brains were extracted, immersed in 30% sucrose, and sectioned into 30 µm coronal slices using a cryostat (Leica, Model 3050S) set to -23 °C. Using the Rat Brain in Stereotaxic Coordinates Atlas (Paxinos and Watson 2009), the regions of interest (ROIs) were localized between -1.9 and -3.36 mm from Bregma, including the CPu, DG, GP, and InsCx. The slices were incubated in PBS at room temperature for 1 hour and then washed three times. Then, they were blocked using a solution of 5% bovine serum albumin and 0.1% Triton X-100 in PBS for 2 hours at room temperature. After blocking, samples were incubated overnight at 4 °C with rat-anti-Iba1 (Ionized calcium-binding adapter molecule 1) antibody (ab283346) to stain microglia and rabbit-anti-NeuN (Neuronal Nuclei) antibody (ABN78) to detect neurons. Next, Alexa 555-conjugated anti-rat (ab150158) and Alexa 647-conjugated anti-rabbit-A647 (ab150079) were added and incubated for 2 hours. The slices were mounted with ProLong® Antifade Kit (Invitrogen, 7481) using 10 - 15 μl per slice. The slides were covered with a coverslip and stored protected from light until further analysis.

### 2.7. Microscopy and data analysis

#### Acquisition

In order to perform cell counts, tile images were taken using a 10x objective on a fluorescence microscope (Zeiss, Axio imager Z1). To obtain detailed images of microglia, microphotographs were acquired using a Zeiss LSM 780 DUO confocal microscope located in the Microscopy Unit of the Institute of Neurobiology and a Zeiss LSM880 AxioObserver confocal microscope from the Microscopy Laboratory of the Center for Applied Physics and Advanced Technology (CFATA). For each sample, twenty Z-stacks were obtained at intervals of 0.8 µm, using a 40x objective and a pinhole of 1 AU.

#### Preprocessing

Confocal images were semi automatically analyzed by region using ImageJ software (v1.54f) and Python. The Iba1 channel was separated and preprocessed, adjusting brightness and contrast, thresholding using the Otsu method, and cleaning with Binary and Noise Remove plugins. A Python algorithm was applied to select and extract cells by detecting areas where the pixels are grouped together and are significantly larger than the smallest pixel groups. At least 50 individual cells per region were saved in separate folders. A visual inspection was conducted to ensure that all selected objects were cells and no other structures or noise. (https://github.com/adebbieeh/Microglia_Morphology_Debbie).

#### Morphology analysis

Morphology of microglia was automatically analyzed by obtaining three main characteristics: number of branches, soma area, and soma perimeter. First, the cells were skeletonized by binarizing the image and tracing the edges from the central part of the soma to the endpoints of the branches. Iteration through the endpoints, slabs, and junctions was used to measure the paths on the skeleton and calculate the number of branches. Additionally, the extraction and measurement of the soma were performed. To do this, small objects, including branches and noise, were removed from the image by defining a specified threshold area, implementing erosion to separate connected objects. Finally, the properties of the resulting objects were used to specifically measure the area and perimeter of the soma.

### 2.8. Statistical Analysis

To identify changes in brain volume a voxel-wise linear mixed-effects model, implemented in R (v.3.6) and RMINC (v.1.5.2.2) software, was employed, utilizing a linear mixed-model GLM with group-by-age interaction (Ctrl vs. Mor) and incorporating subjects as a random effect to capture individual variability. A False Discovery Rate (FDR) correction was applied to correct for multiple testing, with statistical significance set at p ≤ 0.1 (Benjamini and Hochberg 1995). Data on infusions and lever presses were analyzed using linear mixed models (LMMs) with the lmer() function from the lme4 package in R (Bates et al. 2015). The dependent variables included the count of infusions and the total correct and incorrect lever presses. The fixed effects were Group (Ctrl, Mor), Days, and their interaction, with a random intercept for subject identifier (RID) to account for differences between subjects. Post-hoc comparisons were performed using estimated marginal means (EMMeans) from the emmeans package in R (Lenth 2017). Pairwise comparisons between Group levels were conducted for each Day level, and p-values were adjusted using the Tukey method for multiple comparisons. Cell counts and microglia morphology were evaluated for normality and homogeneity of variance using the Shapiro-Wilk test and Levene’s test. When data were normally distributed, either a t-test or two-way Analysis of Variance (ANOVA) was conducted. When homogeneity of variance was not present, Welch’s t-test was used. For non-normally distributed data, the Wilcoxon test was used. To analyze different microglial phenotypes, principal component analysis (PCA) was performed, followed by K-means clustering. The Ctrl and Mor groups were analyzed separately, with the optimal number of clusters determined using the Elbow and Silhouette methods. PCA was then applied to a combined dataset from both groups, and a Fisher’s exact test was conducted to compare cluster distributions between the groups. All analyses were performed in R (v.3.6). Statistical significance was determined at a threshold of p ≤ 0.05.

## 3. Results

### 3.1. Morphine self-administration and drug-seeking behavior in rats

To examine the development of morphine self-administration, we analyzed the pattern of infusions and lever presses. The morphine group showed a progressive increase in the number of infusions along the administration period, with a significant difference observed after the seventh session when compared to the control group exposed to saline solution (F_(19,171)_ = 1.98, p = 0.0116), which did not show escalation in the number of infusions. Following this point, the number of infusions presented intermittent increases, stabilizing during the final four sessions (Fig. 2A). The pattern of total active lever presses increased in a similar manner to that observed for morphine infusions, but with a higher number of lever presses and a difference that approached statistical significance when compared to the control group (F_(19,171)_ = 1.63, p = 0.05), Fig. 2B). In contrast, the total number of inactive lever presses showed a slight increase but did not differ from the control group (F_(19,171)_ = 1.71, p = 0.03. Fig. 2C). Additionally, when analyzing motivation and perseverance combined, we found two rats exhibiting both high motivation and perseverance, and three rats showing only high motivation to obtain the drug (Fig. 2D).

**Figure 2.**
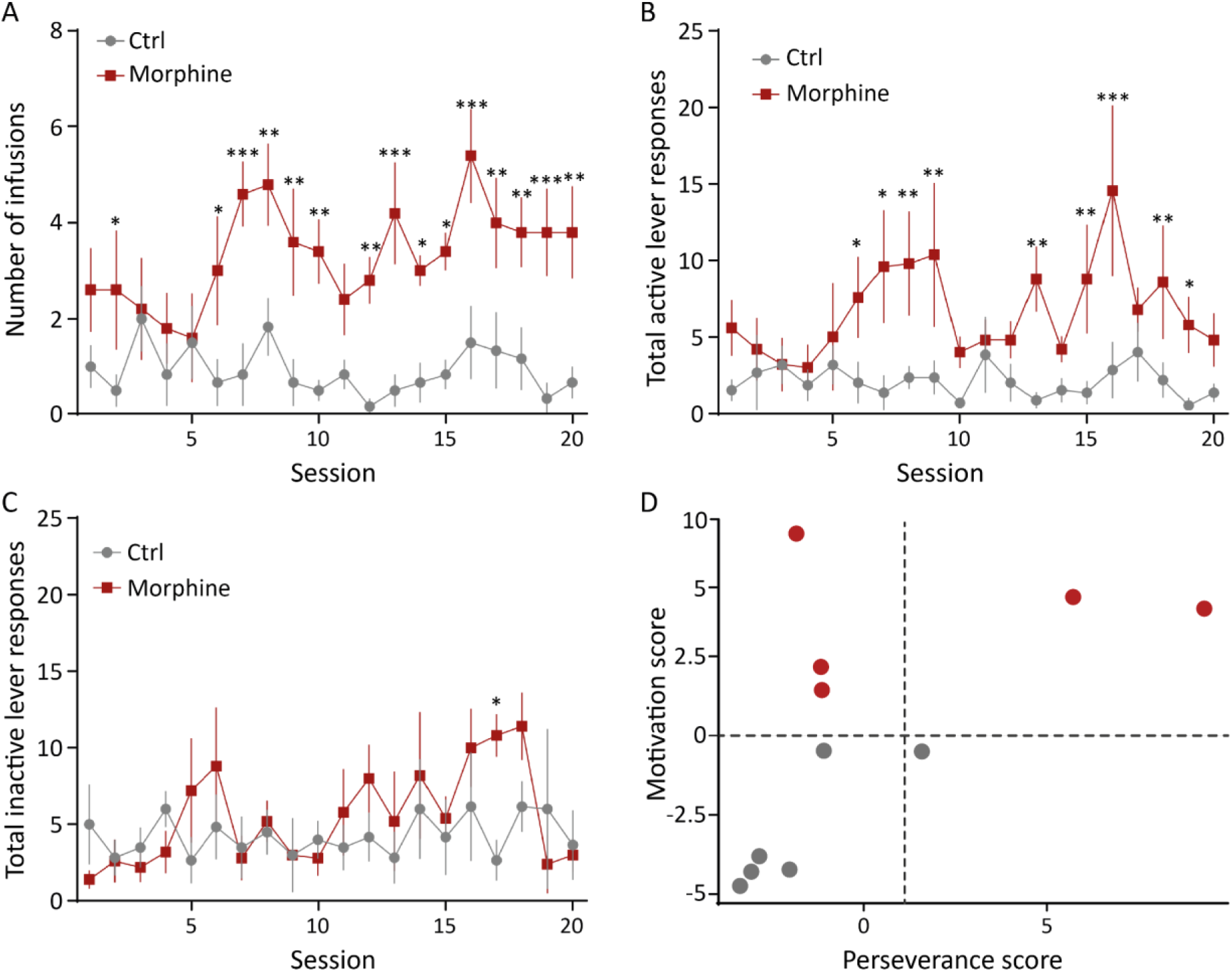
Self-administration behaviors. **A**. Number of infusions per session across a 20-day period. **B**. Total active lever presses per session, including those for obtaining the infusion, during the infusion, and during the time-out period. **C**. Total inactive lever presses, including non-rewarded presses, presses during the infusion, and during the time-out period. **D**. Distribution of control (gray circles) and morphine (red circles) rats, categorized by motivation and perseverance, with individual scores represented. Each data point or square reflects the mean ± standard error of the mean (SEM), with sample sizes of n = 5 for the morphine group and n = 6 for the saline group. *p < 0.05, **p < 0.01, ***p < 0.001, RM-Two-way ANOVA.

### 3.2. Morphine self-administration produces volumetric changes in regions involved in the addictive process

Morphine self-administration modified the local volume in several brain regions (Fig. 3A). In particular, we found an increased volume of a region involving the right globus pallidus (GP) and internal capsule (Fig. 3F, G) and a decrease in the left granular/dysgranular InsCx (Fig. 3J, K), regions involved in addiction processes. Additionally, morphine treatment resulted in an enlargement of the right lobule IX of the cerebellum (Fig. 3B, C), while a reduction in size was observed in the right posterior thalamic nuclear group (Fig. 3H, I). For other regions, changes were expected, for example, an increase in the size of the left pedunculopontine tegmental nucleus (PTg) of the brainstem due to its significant role in opioid modulation of autonomic functions (Gut and Winn 2016). Significant differences compared with control rats are presented through linear regression graphs, highlighting the temporal dynamics and regional impacts of morphine self-administration. Additional regions are shown in table 1.

**Figure 3.**
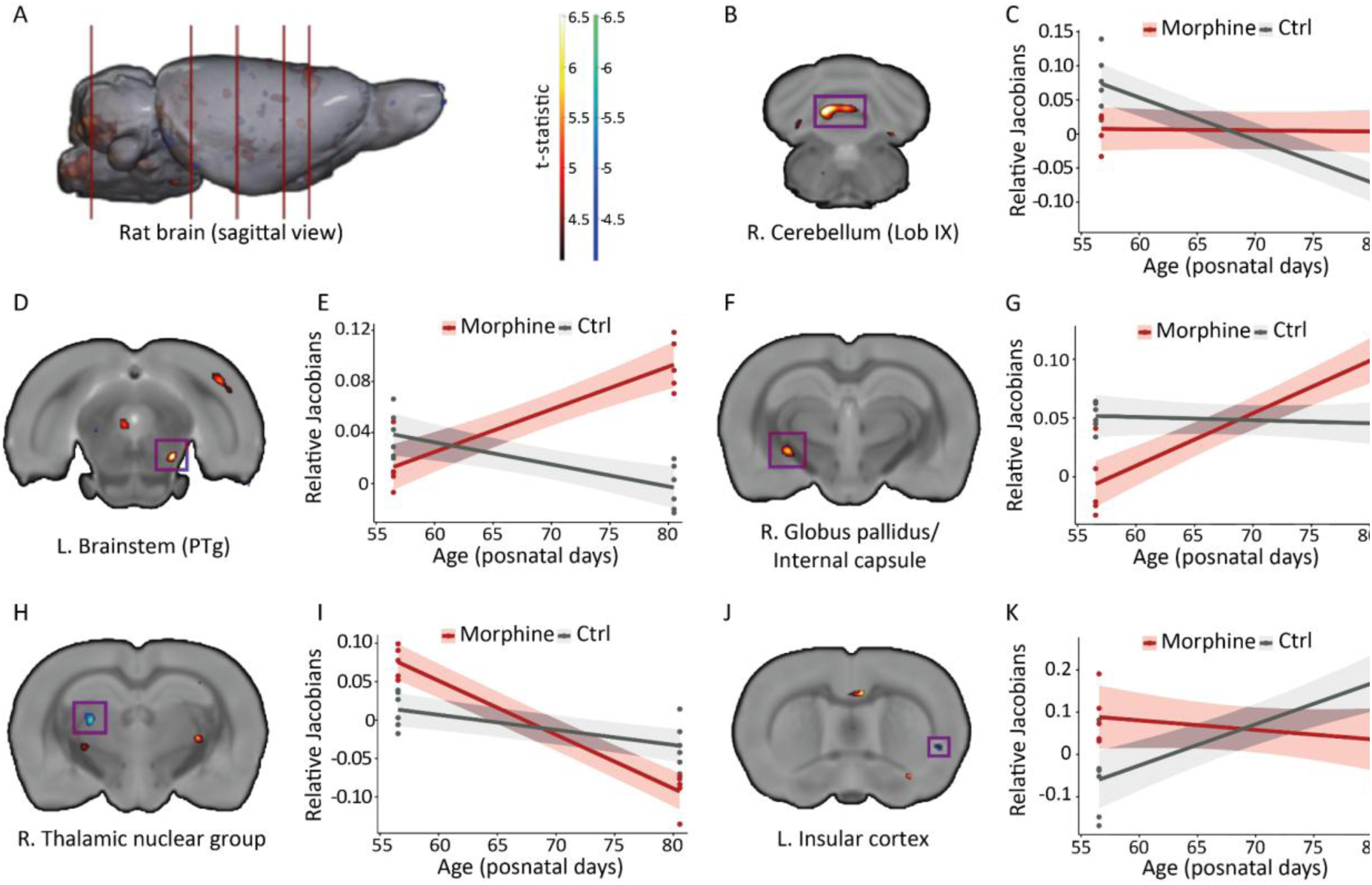
Local volumetric changes obtained through deformation-based morphometry. **A.** Sagittal view of the rat brain indicating the locations of main coronal sections with morphological changes. Coronal sections of the brain with local volume increases (red) or decreases (blue) as well as trajectories of volume (relative Jacobians) with significant changes (morphine vs. ctrl) are shown in regions such as the lobule IX of the cerebellum (Lob IX) (**B, C**), the pedunculopontine tegmental nucleus of the brainstem (PTg) (**D, E**), the globus pallidus/internal capsule (**F, G**), the posterior thalamic nuclear group (**H, I**), and the granular/dysgranular insular cortex (**J, K**). Red lines represent the morphine group, while gray lines represent the control group. R: Right, L: Left. A t-statistic map was used to compare significant differences between the morphine and control groups, with thresholds established using an FDR correction between 10% (t = 4.06) and 1% (t = 6.52).

**Table 1.** Resulting regions of the volumetric analysis.

| ROIs (Paxinos) | Bregma | Sigma Coordinates<br>(x, y, z) | Side | Effect | t-value |
| --- | --- | --- | --- | --- | --- |
| External plexiform layer of the olfactory bulb | 6.60 | 0.46, 7.99, 3.43 | right | Decrease | -5.38 |
| Anterior olfactory nucleus | 5.64 | -1.93, 6.19, 1.93 | left | Increase | 5.08 |
| Primary somatosensory cortex (forelimb) | 1.08 | -4.93, 2.14, 5.08 | left | Increase | 8.80 |
| Primary motor cortex | 1.08 | 2.86, 1.84, 5.68 | right | Increase | 7.36 |
| Layer 3 of cortex | 0.72 | 3.91, 0.79, -2.56 | right | Increase | 4.63 |
| Granular/dysgranular insular cortex | -0.36 | -5.38, 0.04, 0.43 | left | Decrease | -4.33 |
| Paraventricular nucleus of the hypothalamus | -0.72 | -0.28, -0.55, -0.01 | left | Increase | 7.98 |
| Dorsal part of claustrum | -1.32 | -5.53, -1.30, 0.58 | left | Increase | 5.37 |
| Primary somatosensory cortex | -3.36 | -4.03, -2.20, 5.68 | left | Decrease | -4.71 |
| Posterior thalamic nuclear group | -3.72 | 3.01, -3.25, 2.08 | right | Decrease | -9.15 |
| Rostral amygdalopiriform | -3.72 | 5.41, -3.40, -2.71 | right | Decrease | -4.06 |
| Globus pallidus/Internal capsule | -3.72 | 3.91, -3.70, -0.16 | right | Increase | 6.35 |
| Retrosplenial cortex | -5.28 | 0.16, -4.45, 4.18 | right | Increase | 5.74 |
| Pedunculopontine tegmental nucleus | -7.08 | -2.38, -7.00, -0.31 | left | Increase | 7.39 |
| Primary visual cortex, binocular area | -7.32 | -5.23, -7.00, 4.485 | left | Increase | 5.86 |
| Entorhinal cortex | -7.32 | -7.03, -6.859, -1.96 | left | Decrease | -4.16 |
| Sensory root of the trigeminal nerve | -7.92 | 3.01, -8.05, -2.71 | right | Increase | 6.65 |
| Cuneiform nucleus (dorsal part) | -8.76 | -2.08, -8.65, 1.18 | left | Decrease | -6.17 |
| Simplex lobule of cerebellum (SimA) | -9.84 | 3.61, -9.55, 4.18 | right | Increase | 4.23 |
| Paraflocculus | -12.00 | -3.73, -11.80, 0.88 | left | Increase | 5.31 |
| Cerebellar white matter lateral to nucleus VI | -12.00 | 2.11, -11.65, 2.53 | right | Decrease | -4.38 |
| Lobule IX of the cerebellum | -13.36 | 1.51, -14.05, 2.08 | right | Increase | 9.73 |
| Inferior olive of medulla oblongata of the brainstem | -13.36 | 0.46, -14.35, -3.16 | right | Increase | 4.58 |

Regions with significant volumetric changes (group x age interaction) observed in the MRI volumetric analysis (Sigma coordinates). Peak locations are referenced to Bregma according to the Paxinos and Watson atlas (v6). Significance was determined with an $\alpha < 0.05$ and a false discovery rate of $q < 0.05$ . ROI: region of interest.

### 3.3. Cell counts of microglia in regions related to addiction

In order to explore changes at a cellular level, we selected brain regions with volume alterations observed in MRI and relevant literature (Blackwood and Cadet 2021), selecting those most relevant for opioid addiction. Specifically, we selected the GP, a region with a morphine-induced increased volume, and the InsCx, which showed decreased volume in MRI (Fig. 3F, J). Also, we included the CPu and DG, regions involved in the intoxication phase of substance use disorders (Chuang et al. 2011; George F. Koob and Volkow 2016). Neuronal staining was used as an anatomical reference to localize and delimit each region by comparing it with the Paxinos and Watson atlas (2009) (Fig. 4A, B). The density of NeuN-positive cells varied across brain regions, with the DG showing the highest number of neurons, followed by the CPu, InsCx, and GP. However, morphine self-administration did not significantly impact the neuronal count in any of the analyzed regions (Fig. 4C - J, K). In contrast, morphine self-administration resulted in a generalized increase in the number of microglial cells compared with the control group. Specifically, a significant increase in Iba1-positive cells was observed in the CPu, DG, GP, and InsCx (Fig. 4C - J, L), suggesting potential neuroinflammatory responses triggered by morphine exposure in the brain.

**Figure 4.**
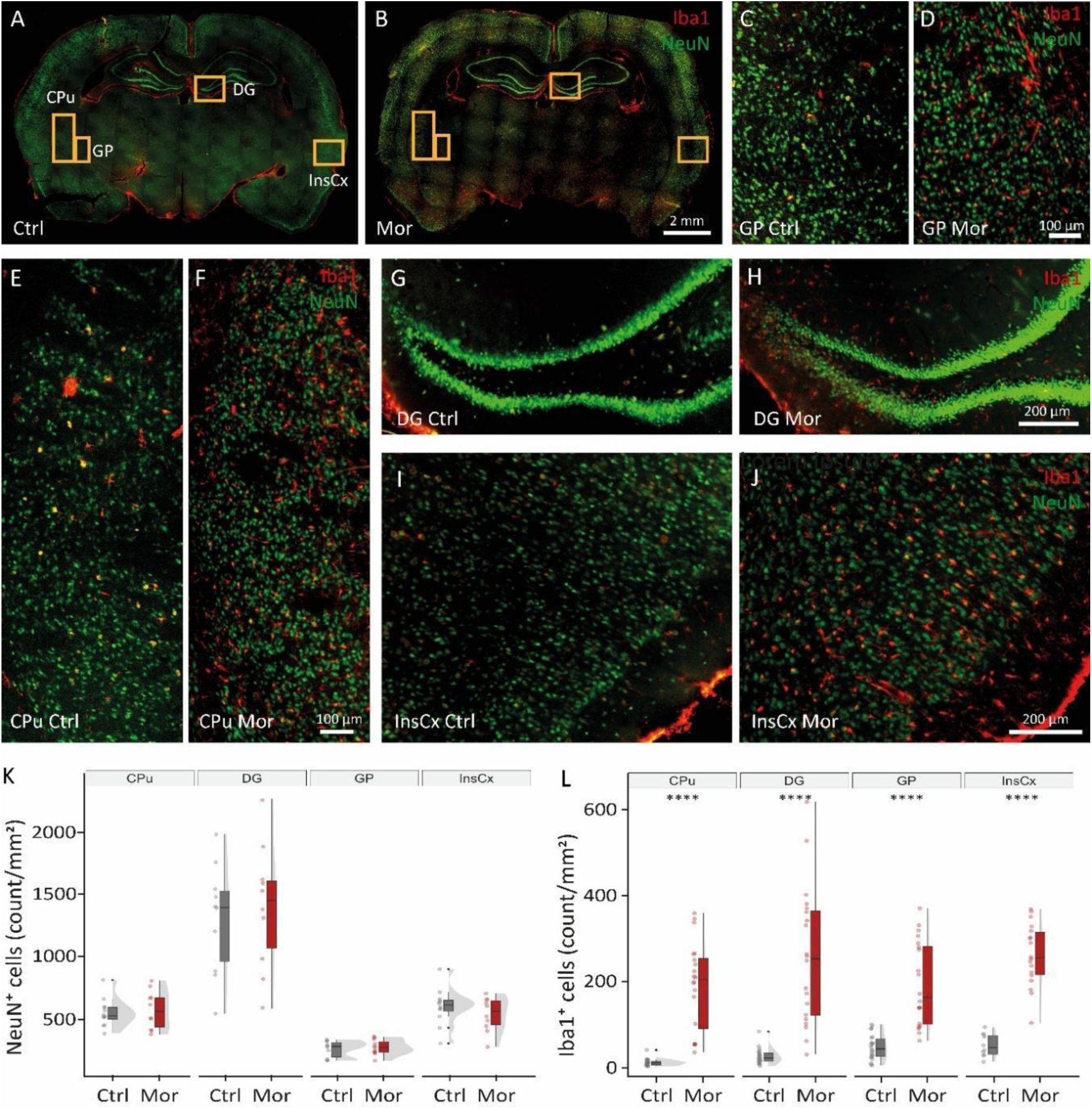
Quantification and distribution of neurons and microglia across brain regions. **A**, **B**. Representative coronal sections from control (Ctrl) and morphine-self-administered (Mor) rats immunolabeled for Iba1 (red, microglia) and NeuN (green, neurons), with regions of interest highlighted in yellow boxes. Magnified views of NeuN^+^ and Iba1^+^ cells are shown for the globus pallidus (GP; **C**, **D**), caudate-putamen (CPu; **E**, **F**), dentate gyrus (DG; **G**, **H**), and insular cortex (InsCx; **I, J**). Quantification of NeuN^+^ and Iba1^+^ cells across the analyzed brain regions (**K**, **L)**. Data are represented as scatter and box plots for both Ctrl and Mor groups. ****p < 0.0001, Wilcoxon test (CPu, DG), t-test (GP, InsCx) (n_slices/animals_ = 9/3). Scale bars: 2 mm (A, B), 100 µm (C, D, E, F), 200 µm (G, H, I, J).

### 3.4. Morphological changes in microglia

Given the observed increase in the number of microglia, we explored different phenotypes of these cells across brain regions. As shown in the diagram in Fig. 5A, cell characteristics were analyzed, including soma perimeter, area, and the number of branches calculated from slabs, junctions, and endpoints. We observed that microglia in the CPu and DG exhibited a significant reduction in both area and perimeter of the soma in the morphine group compared with cells from the control group. In contrast, microglia in the GP demonstrated significant increases in these parameters (Fig. 5B, C). Additionally, we found a lesser number of branches in the morphine group across the CPu, DG, and GP (Fig. 5D). No significant differences were observed in any of the measured parameters within the InsCx when compared to control (Fig. 5B-D). These results suggest that morphine self-administration induces region-specific neuroinflammatory changes, particularly in brain areas associated with substance use disorders.

**Figure 5.**
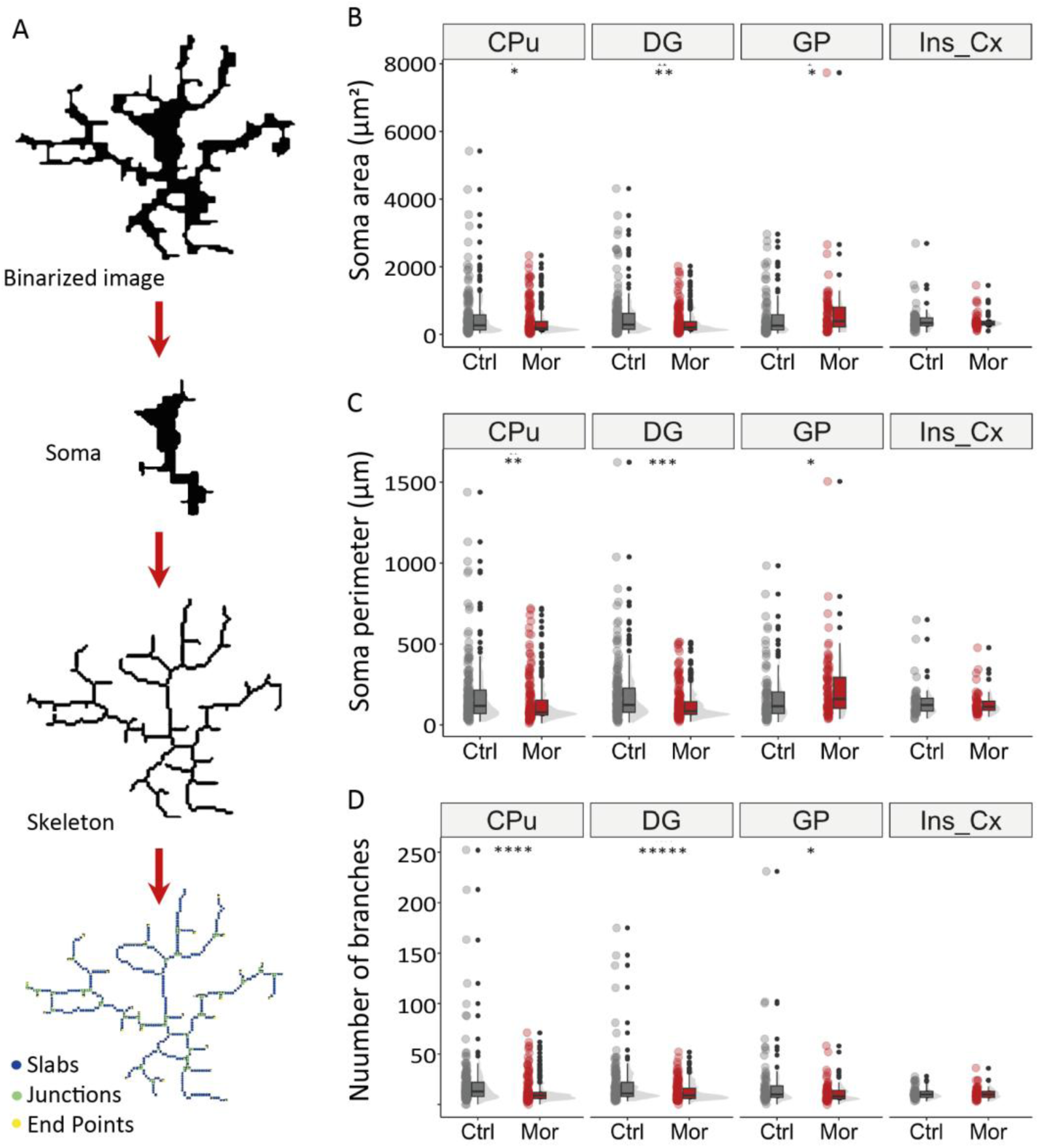
Morphological analysis of microglia. **A**. Representative image illustrating the steps of morphological analysis of microglia including binarization, soma isolation, and skeletonization (slabs, junctions, and endpoints). Quantification of microglial soma area (**B**), soma perimeter (**C**), and number of branches (**D**) across four regions: caudate-putamen (CPu; n = 162 cells per group), dentate gyrus (DG; n = 172 cells per group), globus pallidus (GP; n = 83 cells per group), and insular cortex (InsCx; n = 57 cells per group). Data are represented as scatter and box plots for both Ctrl and Mor groups. *p < 0.05, **p < 0.01, ***p < 0.001, ****p < 0.0001, Wilcoxon test.

### 3.5. Structural characterization of morphine-induced reactive microglia

While individual parameter analysis may offer a general overview of the diversity of microglial phenotypes related to morphine consumption, it may fail to identify important intermediate microglial phenotypes that are essential for understanding their functions. To address this, we systematically identified distinct microglial clusters across various brain regions using the same individual parameters. By PCA and K-means clustering, we enhanced the precision of phenotypic classification. We identified seven clusters in the CPu (Fig. 6A, B), eight in the DG (Fig. 6D, E) and GP (Fig. 6G, H), and six in the InsCx (Fig. 6J, K), highlighting not only the most common microglial phenotypes but also revealing the presence of intermediate morphologies. In the CPu, morphine self-administration led to an increased number of microglia with low branching and larger soma, characteristics typical of an amoeboid phenotype (Fig. 6C, cluster 0). In contrast, fewer cells in the morphine group displayed a moderate number of branches and larger soma, which are indicative of the ramified phenotype (Fig. 6C, cluster 3). In the DG, morphine self-administration increased cells with fewer branches and larger soma, resembling amoeboid-like cells (Fig. 6F, cluster 2). Conversely, it reduced the number of cells with small soma and extensive branching, which are characteristic of hyper-ramified cells (Fig. 6F, cluster 5). Additional clusters of microglia with small somas, long projections, and hyper-ramified phenotypes were absent in the morphine group but present in controls (Fig. 6F, clusters 1, 3, and 6). In the GP, amoeboid and ramified phenotypes were predominant in both groups (Fig. 6I, clusters 0 and 2), with a morphine-induced increase in cells with enlarged soma and abundant branches, resembling a rod-like phenotype (Fig. 6I, cluster 7). In the InsCx, ameboid and ramified phenotypes remained dominant, with no significant differences between the morphine and control groups (Fig. 6L, clusters 0-5). A broad spectrum of other phenotypes was observed across all regions, though without significant changes produced by morphine. These results suggest that while morphine affects specific microglial phenotypes, its influence is region-specific rather than uniform across the brain. Key findings of this project are summarized in Table 2.

**Figure 6.**
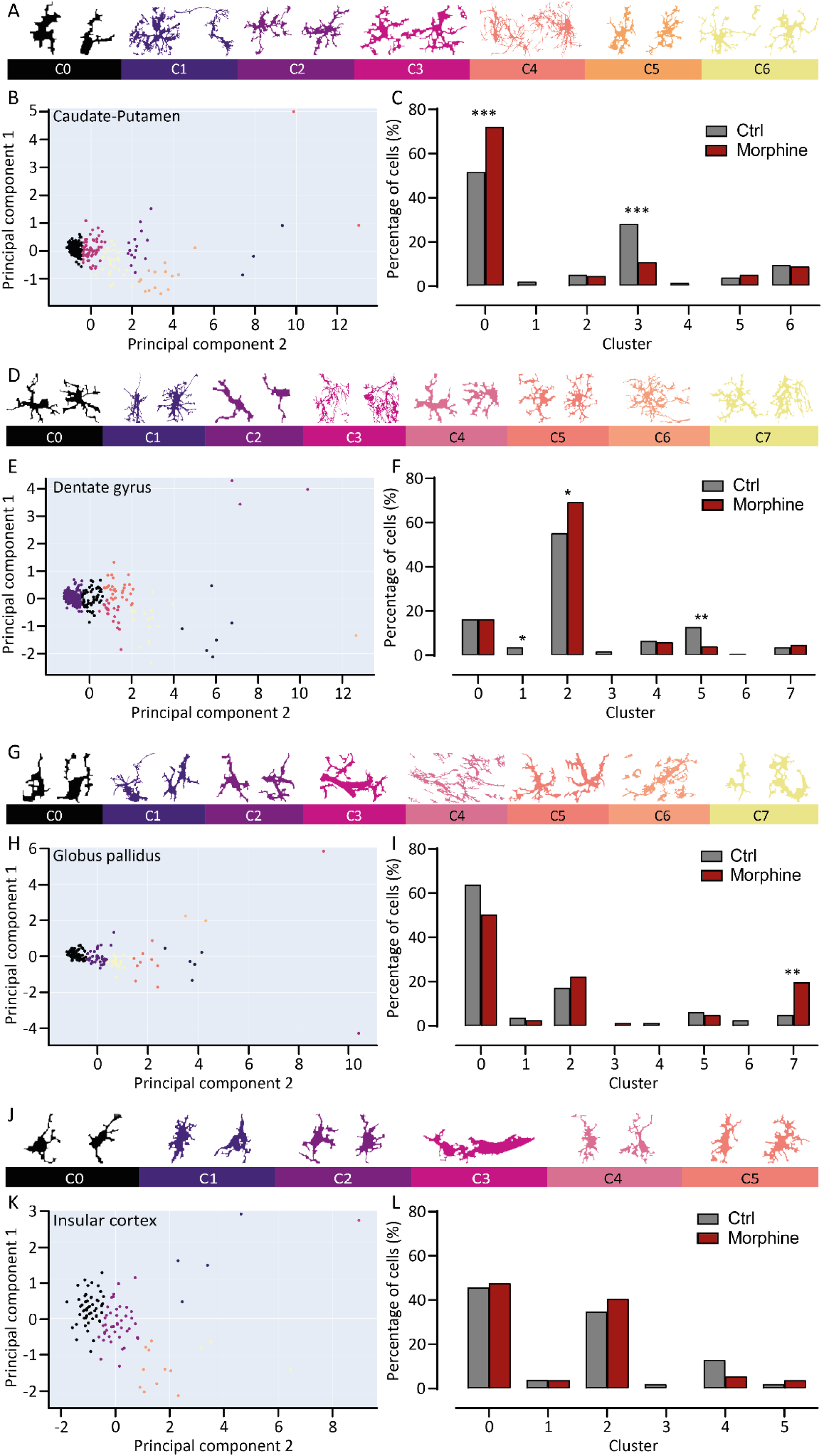
Changes in microglial phenotypes induced by morphine self-administration. Representative 2D images, generated from confocal photomicrographs of Iba1-stained cells using 20 z-stacks per cell, are shown for microglial clusters (C0 to C7) in the caudate-putamen (**A**), dentate gyrus (**D**), globus pallidus (**G**), and insular cortex (**J**). Principal Component Analysis graphs (**B**, **E**, **H**, **K**) illustrating the distribution of total microglial cells, including those from Ctrl and Mor groups in each region. Bar plots (**C**, **F**, **I**, **L**) show the percentage of cells in each cluster, comparing Ctrl and Mor groups. Sample sizes per group: CPu, n = 162 cells; DG, n = 172 cells; GP, n = 83 cells; InsCx, n = 57 cells, obtained from 9 slices from 3 different animals. *p < 0.05, **p < 0.01, ***p < 0.001, Fisher’s exact test.

**Table 2.** Summary of findings.

| ROI | MRI findings | Skeleton morphology | Microglial clusters | Main changes in microglia phenotypes | Proposed biological process |
| --- | --- | --- | --- | --- | --- |
| CPu | No changes | Small soma, fewer branches | 7 | Abundant ameboid cells, fewer ramified cells | Synaptic or extracellular remodeling and neuroinflammation |
| DG | No changes | Small soma, fewer branches | 8 | Abundant ameboid cells, fewer hyper-ramified cells | Synaptic or extracellular remodeling and controlled neuroinflammation |
| GP | Increased volume | Large soma, fewer branches | 8 | Abundant Rod-like cells | Detection of damaged axons, remodeling and neuroprotection |
| InsCx | Decreased volume | Ramified, as in control | 6 | Ameboid and ramified cells, as in controls | Balanced phagocytosis and environmental surveillance |

Summary of key effects of morphine self-administration on brain macrostructure and microglial morphology, including proposed underlying processes. ROI: Region of Interest, CPu: Caudate-Putamen, DG: Dentate Gyrus, GP: Globus pallidus, InsCx: Insular cortex.

## 4. Discussion

Use and misuse of opioids such as morphine have been reported to produce changes in the brain, often linked to adaptive mechanisms that aim to achieve allostasis, which is related with the development of addiction (G. F. Koob and Le Moal 2001). Along with brain alterations, a compulsion to seek and use the drug develops. This behavior is regulated by specific brain regions where opioid receptors, the main targets of morphine, are differentially expressed (Valentino and Volkow 2018; Carranza-Aguilar et al. 2022). Here, we investigated the effects of morphine self-administration on brain macrostructure and cellular microstructure in rats, focusing on the complexity of microglia in regions potentially associated with opioid addiction. In our study we found that chronic morphine self-administration produces volume changes in “classical” brain regions related to addiction (George F. Koob and Volkow 2016) and others such as the cerebellum. It also induces neuroinflammation independent of neuron loss, as evidenced by a higher microglial cell count and alterations in their complexity.

Our findings show that the morphine self-administration protocol we used in rats produced a progressive increase in infusions and a preference for pressing the active lever, indicating drug-seeking behavior (Belin et al. 2013). However, only a subset of rats exhibited both high motivation and perseverance, which contrasts with findings from studies using cocaine, where consistently high scores in both indexes were observed (Bock et al. 2013; Le et al. 2017). This result highlights the individual variability in addiction-like behavior induced by morphine, possibly due to the low dose of morphine we employed. Despite this, the stabilization in the pattern of infusions observed during the final sessions suggests that the rats developed habitual drug consumption, which aligns with findings from other studies demonstrating similar morphine intake patterns in rodents (Kim, Im, and Moon 2018; Zhang and Kong 2017).

To investigate macrostructural changes induced by morphine self-administration, we examined volumetric alterations in the brain and observed changes in the InsCx and GP, both regions crucial for reward processing due to their structural connection with the nucleus accumbens (Cho et al. 2013; Blackwood and Cadet 2021). In the InsCx, we found a morphine-induced volume reduction, a modification consistent with the well-documented effects of opioids on brain morphology in humans (Droutman, Read, and Bechara 2015). Given that the InsCx is important in impulsivity (J. M. Green, Sundman, and Chou 2022) and decision-making processing (Pribut, Sciarillo, and Roesch 2022), diminution of this region may impair these functions, potentially leading to the observed increase in the lever-pressing behavior. In contrast, the increase in the volume of the GP differs from studies in humans reporting decreased brain volume in individuals addicted to heroin (Müller et al. 2019). This discrepancy may be attributed to differences in scan timing and duration of opioid exposure (Tolomeo et al. 2018), since our experiments focused on short-term morphine exposure, whereas human studies typically involve longer periods of drug consumption and postmortem scans (Blackwood and Cadet 2021), where an increase in volume may be related to a more active neuroinflammatory response due to current morphine use. This suggests that the GP is highly susceptible to dynamic structural changes dependent on the duration and intensity of opioid use.

Our study also revealed that morphine self-administration affects brain regions beyond those typically associated with addiction (Table 1). Interestingly, we observed an enlargement of lobule IX in the cerebellum. This region has recently been linked to a circuitry involved in drug-induced conditioning, where increased reactivity to cFOS, a marker of neuronal activity, in lobule IX was observed in rodents exposed to cocaine-associated olfactory cues (Rodríguez-Borillo et al. 2023). Additionally, we observed a reduction in the thalamic volume. In this sense, a study conducted in humans with opioid dependence found decreased functional connectivity between the thalamus and amygdala (Upadhyay et al. 2010), a region involved in motivational withdrawal, which drives compulsive drug-seeking behavior (George F. Koob and Le Moal 2008). The same study reported diminished connectivity between the thalamus and brainstem (Upadhyay et al. 2010). Although alterations in the PTg of the brainstem were expected, the increase we observed differs from other studies that reported a slight reduction in this area in female rats intraperitoneally administered with morphine (Taylor et al. 2023), suggesting a sex-dependent effect of morphine on this region or a self-administration specific change. Further research on the effects of opioids in the PTg of the brainstem is necessary due to its importance in opioid-related pain modulation and autonomic regulation (Mills, Keay, and Henderson 2021; Gut and Winn 2016).

Cellular changes have been proposed to explain volumetric brain alterations in various conditions (Asan et al. 2021). For example, neuronal atrophy could lead to brain volume reductions, while neuroinflammation may contribute to both brain atrophy and volume increases (Datta et al. 2017; Rocca et al. 2017). Since this area of knowledge is poorly explored, we investigated the effects of morphine self-administration on neuronal and microglial counts, focusing on regions with documented volumetric changes (GP and InsCx), as well as areas involved in habit formation and reward-related memories (CPu and DG) (Volkow, Michaelides, and Baler 2019). Our analysis revealed no significant changes in neuronal counts across any of the evaluated regions, indicating that neuronal loss is not a contributing factor to the observed volumetric alterations. In contrast, we found a significant increase in the number of microglia. This increase was observed not only in the GP, where an increased volume was detected, but also in the InsCx, which exhibited a volume reduction. Additionally, the CPu and DG, which did not show any volumetric changes, also displayed an increase in the number of microglia. Therefore, our findings challenge the view that volumetric brain changes directly reflect neuronal loss or gain, and suggest that while some structural changes may be related to neuroinflammation, this mechanism alone cannot fully explain the macrostructural alterations. Other factors that have been reported to affect the macrostructure of the brain include fluid flux, vascular changes, immune cell proliferation, myelin damage, and cell death (Dieleman, Koek, and Hendrikse 2017; Borjini et al. 2019). Additional specific research is needed to better understand these complex processes.

Recognizing that some cellular changes may not be detectable via MRI, and given the evident neuroinflammatory effects of morphine self-administration, we investigated microglial morphology as an indicative measure for specific neuroinflammatory states (Ransohoff and Perry 2009). Analysis of individual microstructural parameters, including soma size and branch count, revealed that morphine self-administration induced a low-ramified microglial phenotype in the CPu, DG, and GP, while the InsCx exhibited a ramified morphology similar to that of controls. In the CPu and DG, the somas were small and rounded, whereas in the GP, they were larger and more elongated compared to controls. Typically, an increase in microglial soma size is associated with phagocytic activity (Davis, Foster, and Thomas 1994), suggesting that these regions are susceptible to damage caused by morphine. However, we did not find changes in the number of neurons, which could indicate other processes associated with this microglial phenotype, such as synaptic remodeling (Weinhard et al. 2018) or extracellular clearance (Nguyen et al. 2020). In the InsCx, a region without observable changes in morphology, morphine-induced alterations in microglia may occur at a slower rate or could be influenced by different neuroinflammatory processes (Ransohoff and Perry 2009).

To better understand the microglial phenotypes induced by morphine, we performed a clustering analysis, revealing increased amoeboid microglia in the CPu and DG. While the CPu also exhibited a reduction in the ramified phenotype, the DG maintained a stable population of non-inflammatory hyper-ramified microglia, suggesting region-specific neuroinflammatory responses. Previous reports indicate that microglia can promote neurogenesis and oligodendrogenesis via pro-inflammatory cytokines (Shigemoto-Mogami et al. 2014). In line with this, our results suggest controlled neuroinflammation in the DG but a more active state in the CPu, suggesting a heightened inflammatory response in this region (Vidal-Itriago et al. 2022). In the GP, where MRI detected increased volume, we observed more rod-like microglia, a phenotype linked to neuroprotection and structural support (Holloway et al. 2019), though *in vitro* evidence suggests these cells may transition to an amoeboid form following morphine exposure (Takayama & Ueda, 2005), shifting from an axon-protective to a phagocytic phenotype in response to morphine. Finally, the InsCx exhibited amoeboid and ramified microglial morphologies comparable to controls, despite a slight reduction in volume. Notably, this region had the lowest number of microglial clusters among the areas studied, suggesting a potential relationship between reduced clustering and the preservation of microglial integrity. This may be linked to sustained microenvironmental modulation, similar to that observed in non-pathological conditions (Hristovska and Pascual 2015).

Limitations of this study include the use of a low dose of morphine, which, while preventing sedative effects, is lower than the doses typically used to induce dependence (Panlilio and Goldberg 2007). Additional research is necessary to explore more clinically relevant doses. Another limitation is the focus on males, as microglial activation varies by sex (Han et al. 2021). Additionally, while morphine affects both opioid and toll-like receptors, our study prioritized analyzing microglial phenotypes to guide future research on underlying mechanisms. Furthermore, we recognize the need to examine microglial responses at different stages of the addiction cycle, such as during abstinence, to determine whether neuroinflammatory changes resolve over time. Future studies will address these aspects to better understand the long-term impact of morphine on the brain.

## 5. Conclusions

In conclusion, our study suggests that brain volumetric changes measured with MRI are not always associated with neuronal loss, as morphine self-administration induces both volumetric and microglial alterations without affecting neuronal counts. This indicates that microglial changes occur independently of neuron loss and may play a role in neuroprotection or synaptic remodeling. The variability in microglial phenotypes across brain regions indicates that multiple mechanisms underlie the effects of morphine, highlighting the complexity of neuroinflammatory responses in addiction. Given that our clustering analysis identified distinct microglial phenotypes, incorporating this approach in future research will be essential for targeting specific brain regions and improving our understanding of how morphine and other substances impact brain function.

## Acknowledgements

This work is partially derived from the bachelor dissertation of Ana Débora Elizarrarás-Herrera from ENES-UNAM Juriquilla. We extend our gratitude to the University Animal Facility Laboratory of the INB for their exceptional technical assistance, particularly José Martín García Servín, María Carbaja Mata and Alejandra Castilla León. MS. Soledad Mendoza Trejo from Lab D12 for the value help on the experimental technical support. We appreciate the invaluable support provided by the Microscopy Unit, including Elsa Nydia Hernández Ríos and Ericka A. de los Ríos Arellano. We also thank the collaborators at the National Materials Characterization Laboratory of the Center for Applied Physics and Advanced Technology (CFATA): Remy Fernand Avila Foucat and Reinher Pimentel-Domínguez. We thank the Behavioral Analysis Unit, Deisy Gasca Martínez. Thanks to Moisés Mendoza from the videoconferencing unit. We are grateful to the National Laboratory for Magnetic Resonance Imaging (LANIREM) for crucial technical assistance, particularly from Juan Órtiz and Luis Concha. Additionally, thanks to M. Mallar Chakravarty and Gabriel A. Devenyi from the Computational Brain Anatomy Lab (CoBrA Lab) (http://cobralab.ca/), CIC, Douglas Research Centre, Montreal, and Compute Canada (www.computecanada.ca). A special thanks to Mallar Chakravarty, Director of the Computational Neuroanatomy Laboratory at Douglas Research Centre, Montreal, Canada, for generously granting us access to computational tools through the Niagara Computing Cluster.

## Authors contribution

EAGV and CJCA designed the study. EAGV provided drugs, reagents and equipment. MACM and CJCA performed the surgeries. ADEH, DMS, MSSR, and CJCA conducted the behavioral experiments. DAV, DMS, MSSR, and CJCA acquired the MRI scans. DAV performed the structural MRI analysis. DMS and CJCA conducted immunofluorescence, image acquisition, and cell counting. ADEH and LATV carried out the morphological and statistical analysis. The draft manuscript was prepared by CJCA, with all authors contributing to and approving the final version.

